# Total immunoglobulin G variation through the phenological cycle of migrant and resident Long-nosed bats *(Leptonycteris yerbabuenae)* in the drylands of Mexico and its relationship with bacterial-killing ability of plasma

**DOI:** 10.1101/2025.06.05.658181

**Authors:** David Alfonso Rivera-Ruiz, José Juan Flores-Martínez, Carlos Rosales, Luisa I. Falcón, Marco Tulio Solano de la Cruz, L. Gerardo Herrera M.

## Abstract

Immunological variations of bats throughout their life cycle represent an underexplored and controversial topic that has great potential for understanding their immunology. This gap is particularly large regarding the state of immunity during early life and migration, and how different immune components interact with each other. In this study, the partial migratory long-nosed bat (*Leptonycteris yerbabuenae*) was used as a model to assess the state of acquired humoral immunity (total IgG concentration, tIgG) during early life, during the reproductive cycle and at the extremes of the migratory range of females. We also determined the relationship of tIgG with plasma bactericidal activity against *Escherichia coli*. The concentration of tIgG was stable throughout the reproductive cycle of adults, and no significant changes were detected at the extremes of the migratory range of females. However, tIgG concentration was reduced during early life (2 to 3 months of age) and in non-reproductive males in March-May 2019 relative to non-reproductive males in March 2020. The tIgG concentration had a positive relationship with plasma bactericidal ability in non-reproductive females, but no relationship was observed in non-reproductive males and reproductive females. These results suggests that tIgG concentration is resilient to physiological and ecological changes experienced by adults throughout their reproductive and migratory cycle. The reduction of tIgG during early life suggests that young individuals have not developed an IgG pool comparable to that of adults. Finally, the positive relationship observed between tIgG and BKA in non-reproductive females suggests that *E. coli* clearance is enhanced by the presence of antibodies that facilitate the elimination of this bacteria.

## Introduction

Mammalian immunity comprises innate and acquired defense mechanisms that work in synergy to control different types of micro-organisms and parasites that colonize their bodies [1,2]. Both branches of the immune system modulate their response depending on the stimulus detected [3,4] and are closely linked to the energetic, hormonal and environmental condition of the individual [5–8]. This immune plasticity may be associated with demographic factors such as age [9], reproductive activity [10] and migratory movements [11]. In bats, the relationship between immunity and demography has been explored in a small number of species despite their high biological diversity [12] and immunological plasticity [13,14]. Bats have unique immunological characteristics that allow them to coexist with micro-organisms of great health importance [15–18], whose life cycles can vary throughout the life cycle of their hosts [19–21]. Therefore, it is essential to determine the immunological changes that bats undergo during their life cycle to identify possible periods of increased likelihood of infection.

The immune system of wildlife mammals during early life has been little explored. The ontogenetic changes that an individual undergoes from early life to adulthood are mainly related to acquired immunity [9,22–24], although some aspects of innate immunity differ between young and adults [25]. In bats, young are more susceptible to infectious agents, which increase the rate of infection within the population and the possibility of zoonotic outbreaks [19,26]. Despite this increased susceptibility during early life, bats manage to coexist with these threats, but the performance of their immune system during early life has been scarcely studied. Like other mammals, young bats acquire maternal antibodies that decline after weaning [27–30], which suggest that humoral acquired immunity is not fully developed at this stage [31]. In contrast, other immunological parameters are fully developed during early life as is the case for phagocytosis and lymphocyte proliferation [32].

Reproduction affects many aspects of mammalian biology that can influence immune function [33–35]. In bats, the immune system may be altered by the increased risk of infection [19,36–38] and the hormonal [39], microbial[40–42] and redox [43] changes experienced by reproductive individuals. For example, white blood cell (WBC) counts increase in reproductive males of *M. daubentonii* but total immunoglobulin G (tIgG), hemolysis, hemagglutination and bacterial-killing ability (BKA) are not affected [44]. In contrast, reproduction in males of *Desmodus rotundus* enhance BKA but decrease plasma tIgG, which suggests an immune trade-off between innate and acquired immunity during reproduction [45]. The inflammatory response to phytohemagglutinin (PHA) may also be altered during reproduction in males of *M. vivesi* [39] and *L. yerbabuenae* [46]. In female bats, tIgG and hemolysis increase during pregnancy and lactation in *M. daubentonii*, respectively, but WBC count, hemagglutination and BKA did not vary between reproductive stages [44]. In contrast, BKA in *M. vivesi* [39] and the inflammatory response in *M. myotis* [36] and *M. vivesi* [39] during gestation.

Migration is a versatile movement strategy that, in most species, is associated with periodic movements that seek to exploit the most energetically favorable conditions for reproduction in seasonal climatic environments [47,48]. Migratory movements might affect immunity by either increasing [49–51] or reducing the risk of infection [52–54], and by the energy costs [55] and oxidative stress [56] associated with this behavior. The effect of migration on the immune system of bats has only been studied in a handful of species. Migration does not affect basal haptoglobin levels of *Pipistrellus nathusii* but affects the number of systemic lymphocytes and neutrophils [57]. Bacterial lipopolysaccharide (LPS)-induced immune response increases haptoglobin levels only during the active phase of migration in *P. nathusii* while this immunological challenge does not affect the number of systemic lymphocytes and neutrophils [57]. Males of *Lasionycteris noctivagants* that migrate long-distances show no immunological changes respect resident males, in contrast to males that migrate short-distances, which show reduced tIgG levels when they come from the north or a higher N:L ratio and higher neutrophil numbers when they come from southeastern regions [58].

We have previously tested the effects of demographic factors on the innate immune response of the long-nosed bat (*Leptonycteris yerbabuenae* (Phyllostomidae)). This bat is mainly distributed in the arid and semi-arid regions of Mexico [59]. Most females in the Pacific coast mate in winter in central-western Mexico [60,61] and migrate to higher latitudes to form maternity colonies in spring-summer [62–64]. In contrast, males are resident through the year [65–69]. We found that migratory females that have just arrived in the central-western region of Mexico have lower plasma BKA compared to females captured two months later in the same region and to pregnant migratory females from the northern region of Mexico, that young do not exhibit a BKA difference from that of adult females captured in the same region, and that reproductive males exhibit a higher BKA than non-reproductive males [70].

Here we extended our work with *L. yerbabuenae* to test if the humoral acquired immunity varied with demographic factors. We explored the relationship of age, migration and reproductive status on tIgG in *L. yerbabuenae*. We hypothesized that young should have lower tIgG than adults, as the concentration of this antibody is reduced during early life [71–75]. We expected that levels of tIgG may be reduced during the male reproductive season because reproduction [45] and testosterone [10,76] may negatively affect tIgG production. The only evidence reported in bats suggests that gestation increases tIgG [44]; therefore, we propose that this same pattern is likely to be observed in *L. yerbabuenae*, especially if pregnant migratory females may still be dealing with potential microorganisms and infectious agents present along their migratory route [49,77]. We predicted that lactating females should not exhibit changes in tIgG as in *M. daubentonii* [44] and because it is unlikely that the effect of migration can still be reflected in the performance of in their immune system as these females arrived in the region 2 to 3 months before sample collection. Migratory females returning to west-central Mexico could exhibit a higher concentration of tIgG if they were exposed to greater diversity of infectious agents during their return journey from higher latitudes [49,77]. We also examined the relationship of tIgG and BKA in non-reproductive adult males and females, and reproductive females, and expected a positive correlation because some IgG isotypes are potent activators of the complement system[78], which is the main immunological component associated with BKA [79]. However, it is also possible that an antagonistic relationship between tIgG and BKA may be observed as is the case in *D. rotundus* [45].

## Materials and Methods

### Study sites and sample collection

We collected samples of *L. yerbabuenae* in west-central Mexico in Don Panchito Island, Chamela, Jalisco (19°32’08.4’’N, 105°05’18.8’’W) and in the Fábrica cave in Coquimatlán, Colima (19°09’05.8’’N, 103°50’06.9’’O), and in northwestern Mexico in the Mariana cave between Carbó and San Miguel de Horcasitas, Sonora (29°35’25.9’’N, 110°48’8.9’’W; Fig Adul females and reproductive and non-reproductive adult males were captured in Chamela and Coquimatlán in March, May, October and December of 2019, and in Chamela in March 2020. In October-January the colony in Don Panchito reaches its maximum size due to the arrival of migratory females in synchrony with the reproductive activity of males [60,61,80]. After this period, many adult females progressively disappear from the colony, which is predominantly composed of non-reproductive adult males from February to August [60,61]. Although the population dynamics in Coquimatlán has not been described, it is probable that it follows the same pattern as in Chamela. It has been suggested that most females migrate towards higher latitudes in spring and summer to give birth, following the blooming of columnar cacti and agaves [62,81,82]. This hypothetical migratory route is supported by the fact that young individuals, and lactating and pregnant females appear during May-August in northwestern Mexico and southwestern USA [62]. Accordingly, we collected samples of young and pregnant and lactating females in the Mariana cave in May and July-August 2019.

The Ethics Committee in Research and Teaching of the Institute of Biology, National Autonomous University of Mexico approved the protocol of the study. We followed the guidelines established in the research permit authorized from Dirección General de Vida Silvestre (#8053/199). The capture and sampling of bats were carried out in accordance with the guidelines of the American Society of Mammalogists [83]. No animal was subject to anesthesia or euthanasia. Bats were captured between 22:00 and 06:00 hours with mist nets placed near the entrance of the Mariana cave, and with a hand net inside the caves in Chamela and Coquimatlán. Each individual was placed in a clean cotton bag until processing. Blood samples (ca. 100 µl) were collected within 30 minutes after capture by antebrachial vein puncture with a needle (30G x 1/2’’) using a micropipette and sterile tips previously treated with heparin (Inhepar®, 5000Ul/ml). Blood was placed in sterile heparin-impregnated Eppendorf tubes and centrifuged at 6000 rpm for 4 minutes to separate plasma from blood cells. Blood plasma was collected in cryotubes and placed in liquid nitrogen for transport and final disposal at -70°C in laboratory. Finally, we recorded the sex, age category (adult or young), reproductive condition, length forearm (Mitutoyo CD-6, Mexico; ±0.01 mm) and body mass (Ohaus, Nueva Jersey, USA, ± 0.1 g) of each individual. Young should be approximately 2 to 3 months old as they were captured in July, and pregnant migratory females arrive in the Sonora region around May to rear their young [62]. We distinguish adults from young by examining multiple phenotypical traits: the classical epiphyseal– diaphyseal fusion of the 4th metacarpal–phalangeal joint [84,85], the hair and the body mass index (BMI; see below). In contrast to adult females in Sonora, young individuals had short grayish hair, and wide epiphalangeal sutures (in almost all phalanges). Reproductively active males of *L. yerbabuenae* are characterized by scrotal testicles and a prominent patch on the scapular region, which is involved in attracting females [86,87]. Although there are males with scrotal testicles during April-July in Don Panchito Island, only the September-December males with scrotal testicles and the dorsal patch produce semen [80]. For females, we classified them as pregnant by palpation of their abdomen and lactating if they presented hairless nipples that released milk after being pressed. Females captured between October and December on Don Panchito Island are probably mating but do not present external morphological characteristics that indicate their reproductive status and were arbitrarily classified as non-reproductive.

### Body mass index (BMI)

Previous studies have determined that fat stores of *L. yerbabuenae* on Don Panchito Island vary throughout the year [61]. Therefore, we calculated the body mass index (BMI: body mass/ forearm length) as a measure of body condition across different demographic cohorts [88–92]. This ratio is intended to determine the magnitude of fat reserves as a function of individual size, as individuals with a larger lean mass should be heavier than smaller individuals regardless of accumulated fat reserves [93].

### Bacterial-Killing Ability (BKA) of plasma

Most of the BKA data were acquired from Rivera et al (2023) [70] and complemented with additional samples that were not included in the original article. The BKA of these additional samples are presented here as supporting information (S1 Fig. and S2 Fig.). The experimental design of plasma BKA was based on the protocols proposed by Liebl and Martin [94] and French and Neuman-Lee [95], in which the *Escherichia coli* strain ATCC 8739 (Microbiologics, St. Cloud, MN, USA) was used as a target microorganism to quantify the ability of bat plasma (natural antibodies, complement system and lysozyme) to inhibit its growth. The BKA assay is a common immunological measure in bat studies [13,39,43–45,79,96–102]. To avoid contamination, experiments were conducted in a laminar flow hood using autoclaved equipment and sterile disposable materials. Lyophilized pellets of *E. coli* were reconstituted in a phosphate-buffered solution (PBS) and diluted to a working concentration of 10^5^ bacteria/mL. Plasma samples were thawed only once, and they were used immediately. We used 96-well conical-bottom cell culture microplates (Sarstedt, Nümbrecht, Germany, 83.3926.500) to plate 9 replicates of positive controls (18 µL of 0.01 M sterile PBS; Sigma Aldrich, St. Louis, MO, USA), 6 replicates of negative controls (18 µL of PBS), 9 replicates of an antibiotic control (2 µL of ampicillin at a concentration of 0.05 µg/mL in 16 µL of PBS), and 3 replicates of each sample (2 µL of plasma diluted in 16 µL of PBS). We then added 4 µL of a bacterial suspension with a concentration of 10^5^ bacteria/mL to all wells except the negative controls. The plate was partially covered during all procedures that required removal from the cabinet to avoid contamination (incubation and absorbance measurement) and to allow oxygen flow. After the plate was covered, the plate was incubated at 200 rpm and 37 °C for 30 min. Later, 128 µL of nutrient broth (BD 234,000) was added to each well, and the absorbance of the plate was recorded at a wavelength of 600 nm (Epoch Microplate Reader, Biotek, Winooski, VT, USA). Following this measurement, the plate was incubated for a second time at 200 rpm for 12 h at 37 °C. Then, we measured the absorbance at 600 nm (Epoch Microplate Reader, Biotek). The negative control was used to ensure that there was no contamination, and the antibiotic control was used to verify that the bactericidal action could be evaluated with the assay, but they were not used in the final calculation of BKA. The calculation of plasma BKA was determined as the percentage of bacteria eliminated by the plasma in relation to the positive control, using the mean of the replicates for each parameter, with the following formula:

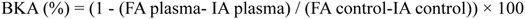

where:

1) FA plasma = Final plasma absorbance
2) IA plasma = Initial plasma absorbance
3) FA control = Final absorbance of the positive control
4) IA control = Initial absorbance of the positive control

### Quantification of total immunoglobulin G plasma concentration

As a measure of acquired humoral immunity, we quantified total immunoglobulin G concentration (tIgG) by enzyme-linked immunosorbent assay in 96-well plates (SARSTEDT "ELISA-Plate, High Binding 82.1581.200). Because plasma tIgG had not been measured previously in *L. yerbabuenae*, we performed a pilot test to determine the ideal plasma dilution. We randomly selected two samples (2µL per sample) representing non-reproductive and reproductive males, non-reproductive and reproductive females, and young individuals. The plasma samples were subjected to double serial dilutions using sodium bicarbonate as solvent (NaHCO_3_, 0.1M, pH= 9.6), within a dilution range of 1/625 (2 µL of plasma in 1248 µL of NaHCO_3_) to a dilution of 1/80000. Each of the dilutions performed were worked in triplicate, where a volume of 50 µL per well was used.

The plate with the plasma dilutions was incubated for 18 hours at 4°C to favor the binding of the plasma proteins to the bottom of the wells. Following this incubation, most of the diluted plasma volume was removed and 2 washes were performed with 200 µl of 0.05% PBS-Tween-20 to remove excess sample. The next step consisted of blocking the bottom of the plate with 50 µl of bovine serum albumin (BSA) diluted 1% in PBS 1X. The plate was incubated for 2 hours at room temperature and then washed 2 times with 200 µl of PBS-Tween-20 0.05%. To detect plasma tIgG, a volume of 50 µL of peroxidase-coupled anti-IgG antibody (Goat anti-Bat IgG (H+L) Secondary Antibody [HRP], NB7238, NOVUS BIOLOGICAL) was added at a 1/10000 dilution and allowed to react for 2 hours at room temperature. After incubation, two washes were performed with 200 µl of PBS-Tween-20 0.05% and 50 µL of SIGMA FAST-OPD (p9187) developer solution were added to each well, and the colorimetric reaction was stopped after 5 minutes with 100µl of H_2_SO_4_. Finally, the absorbance of the plate was recorded from 450 to 500 nm to detect the most appropriate wavelength.

The average absorbance of the triplicates of each of the 10 serially diluted samples was calculated to determine the most appropriate dilution. From the range of dilutions in which tIgG decreased as the dilution factor increased (S3 Fig. and S1Table), the 1/20000 dilution was determined to be the most appropriate as it has the steepest slope (highest rate of change per dilution factor). The absorbance was recorded at 490 nm because the highest absorbance was observed at this wavelength. Therefore, all remaining samples were worked in triplicate and diluted at a ratio of 1 part plasma (2 µL) to 20000 NaHCO_3_ using the same procedure used for assay standardization. Antibody concentration for each sample was determined from the average optical density (OD) of three replicates, as color intensity is directly proportional to antibody concentration.

### Statistical analysis

R software version 4.3.2 was used for statistical analysis [103]. According to the results of the fitdistrplus package [104], the lognormal and normal probability distribution functions were the most appropriate for the analyses of BMI and tIgG, respectively. The analyses of both variables were performed with a general linear model and the statistical significance was determined by means of a deviance analysis using the ANOVA function of the car package [105]. When an explanatory variable significantly affected the response variable, the multcomp package was used to perform a Tukey multiple comparisons test to identify cohorts that differed from each other [106] and for the case of interacting factors we used the emmeans package [107]. The statistical models used to assess demographic variation in tIgG did not include BMI as a covariate because body condition was not homogeneous within the demographic groups (Fig 2), which violates the principle of independence of explanatory variables [108]. Therefore, BMI and tIgG were evaluated by independent models where the demographic factors were considered as explanatory variables.

**Fig 1.**
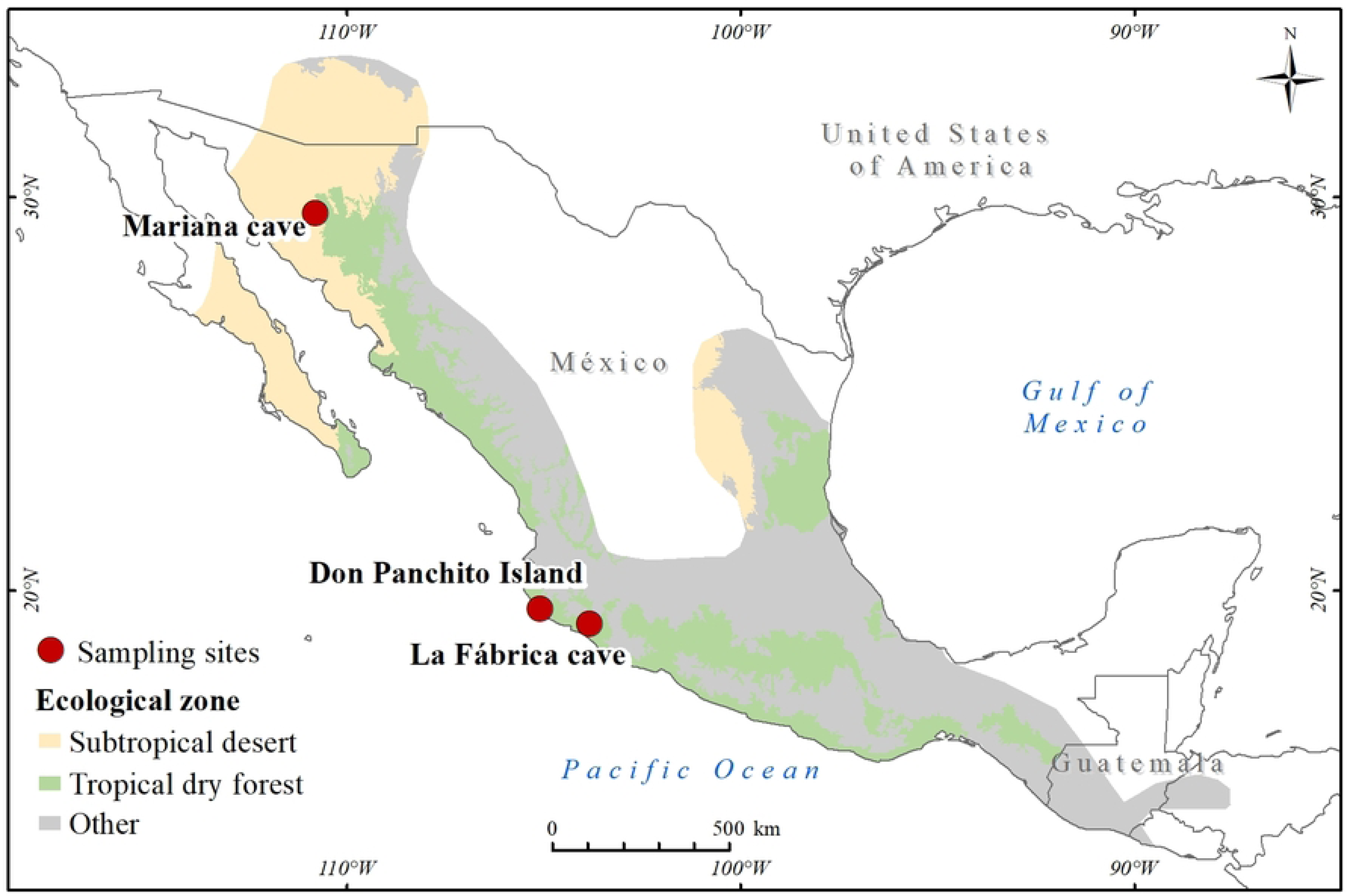
Geographical range and main vegetation types of the ecological zones inhabited by the Lesser long-nosed bat (*Leptonycteris yerbabuenae*) in Mexico. The range distribution data were acquired from the IUCN red list of threatened species [59]. The subtropical desert (beige) and the tropical dry forest (green) represent the main ecological zones occupied by *L. yerbabuenae* (other ecological zones are indicated in gray in the category ‘*Other*’). The red dots indicate capture and sampling sites. during 2019 -2020.

**Fig 2.**
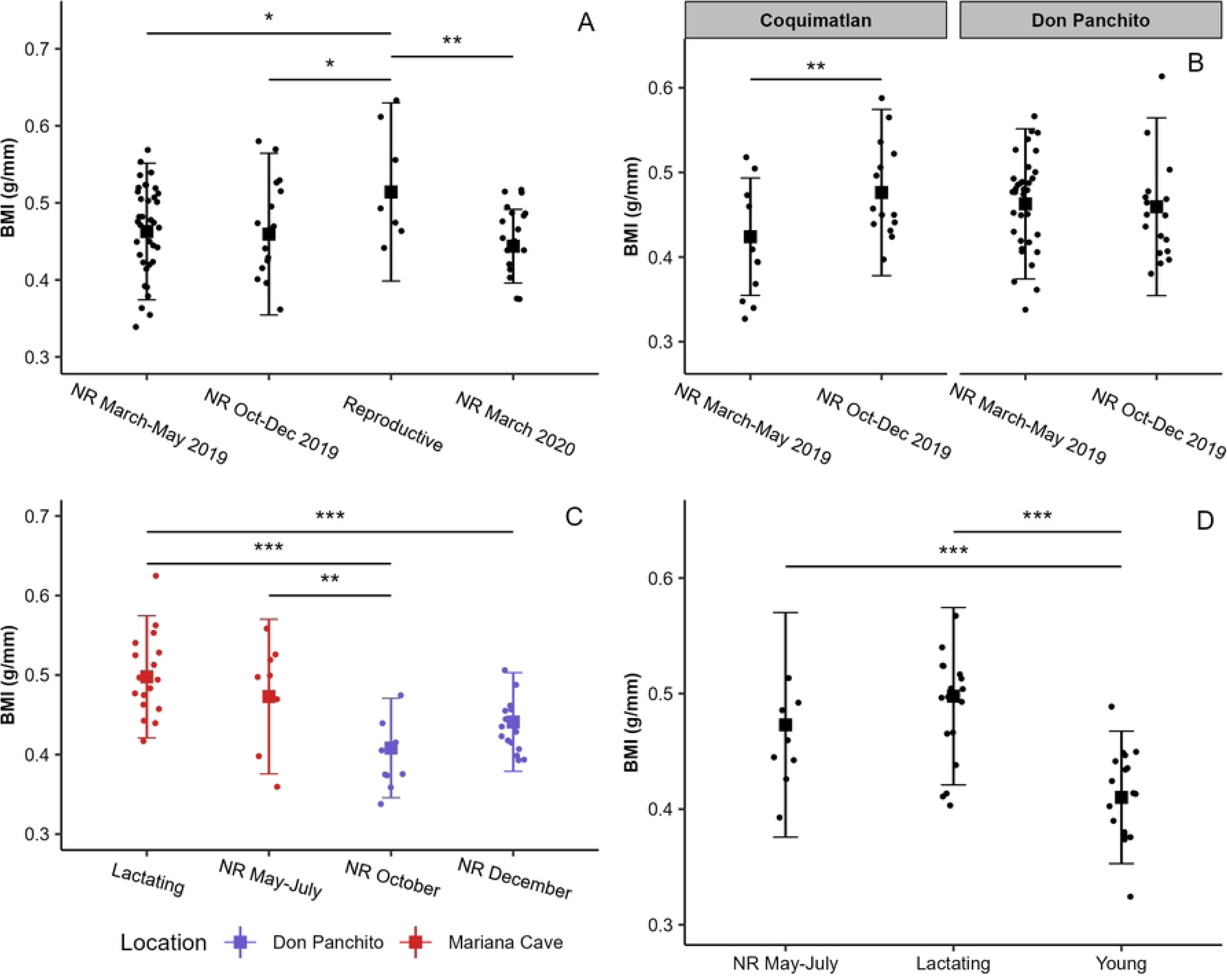
Body mass index (BMI) across different demographic cohorts of the Lesser long-nosed bat (Leptonycteris yerbabuenae) in Mexico. A) reproductive and non-reproductive males in Chamela during March and May (NR March-May 2019) and October and December (NR Oct-Dec 2019) in 2019, and in March 2020 (NR March 2020), B) non-reproductive (NR) males of Coquimatlán and Chamela during the months of March and May (NR March-May) and October and December (NR Oct-Dec) in 2019, C) lactating and non-reproductive females of Sonora (NR May-July), and non-reproductive females in October (NR October) and December (NR December) in Chamela, D) lactating, non-reproductive adult females (NR May-July) and young’s in Sonora.

The effect of male reproductive activity on tIgG it was determined by comparing the reproductive cohorts in December with the non-reproductive cohorts in March-May and October-December 2019 and March 2020 in Chamela. To evaluate if tIgG differed between non-reproductive males from Coquimatlán and Chamela in March-May 2019 and October-December 2019, we used a statistical model that considered location and season as main factors and their interaction. Since the different reproductive phases of females occur concomitantly with migration, it is not possible to discern the isolated effect of both factors; therefore, pregnant and lactating migratory females from Sonora (May and July 2019) were compared with non-reproductive migratory females from Jalisco (October-December 2019) and Sonora (March-May 2019). To address the degree of development of acquired immunity during early life stages, young from Sonora were compared with lactating and non-reproductive females that were captured in the same region and time (May and July 2019). Sex can affect bat immunity [39,43,44,101], and although it is possible to assess the effect of this variable between non-reproductive cohorts of males and females of Jalisco in October-December 2019, we discarded this comparison because females had probably completed migration, which would obscure physiological differences associated with sex. BMI analysis was performed across the same demographic groups used for tIgG. Outliers that could significantly affect the models for BMI (we removed 2 non-reproductive males of Chamela) and tIgG (we removed one non-reproductive female and one young) were identified by means of Cook’s distances. Finally, all models were validated by residual analysis using the DHARMa package [109].

We tested the relationship between BKA and tIgG with a Spearman correlation tests because BKA had a non-parametric distribution with many tied data (many individuals have 100% and 0% values) [110] for each of the following cohorts: non-reproductive females, reproductive females (we pooled pregnant and lactating females because there was no immunological difference between them), and non-reproductive pooled males of Coquimatlán and Chamela (none immunological differences were detected between these cohorts). The immunological parameters of young and reproductive males were not correlated due to the low sample size. Data is presented as mean ± SD along the text.

## Results

### Body mass index varied seasonally and demographically

Only two reproductively active males were captured in Coquimatlán during the October-December 2019 season, which were not included in the analysis. In Chamela, reproductive males were captured in December 2019 (n = 7), and non-reproductive males in March-May (n=37) and October-December (n=19) 2019 and March 2020 (n=19). We found significant evidence that BMI differed among all categories (F_3,76_ = 3.99, p=0.01; n=80, Figure 2A): reproductive males had marginally higher values than non-reproductive males in March-May (p=0.04) and October-December (p=0.03) 2019, and that in March 2020 (p=0.004). In contrast, BMI was not affected by locality (F_1,76_ = 1.39, p = 0.24, n=80, Figure 2B) and season (F_1,76_ = 2.19, p = 0.14, n = 80, Figure 2B) in non-reproductive males of Chamela and Coquimatlán. However, we detect a significant effect of the interaction between locality and season (F_1,76_ = 6.39, p = 0.01, n = 80, Figure 2B), which is driven by a lower but significant increase in BMI (p = 0.004) in males from October-December 2019 (0.47 g/mm ± 0.04) in Coquimatlán respect of males in March-May 2019 (0.42 g/mm ± 0.03). In contrast, non-reproductive males in Chamela did not exhibit seasonal variations in BMI (p = 0.76).

Pregnant females from Sonora and one lactating female from Chamela were excluded from the BMI analysis due to additional fetal weight and low sample size, respectively. Lactating (n=17) and non-reproductive (n=8) females from Sonora (May-July 2019) presented differences in BMI from their conspecifics captured in October (n=9) and December (n=17) 2019 in Chamela (F_3,47_ = 14.08, p < 0.001, n=51; Figure 2C). Non-reproductive females from Chamela captured in October had lower BMI than lactating (p < 0.001) and non-reproductive females from Sonora (p = 0.002), while females in December 2019 had lower BMI than lactating females from Sonora (p < 0.001). Finally, young in the Sonora region (n=17) had lower BMI than lactating females (n=17) and non-reproductive females (n=8) captured in the same period in Sonora (F_2,39_ = 25.80, p < 0.001, n=42; Figure 2D).

### The concentration of tIgG across different demographic cohorts

The tIgG concentration varied seasonally in non-reproductive males from Chamela (F_3,78_ = 4.08, p = 0.009, n=82; Figure 3A), with higher values in March-May 2019 than in March 2020 (p = 0.004). However, the tIgG concentration of reproductive males did not differ from that in males in March-May (p = 0.51) and October-December (p = 0.96) 2019, and March 2020 (p = 0.78). Additionally, tIgG concentration of males in October-December 2019 did not differ from those in March-May 2019 (p = 0.59) and March 2020 (p = 0.22). There was no difference of tIgG concentration between non-reproductive males of Coquimatlán and Chamela (F_1,78_ = 0.37, p = 0.54, n=82; Figure 3B), and season (F_1,78_ = 0.19, p = 0.66, n=82; Figure 3B) and its interaction with locality (F_1,78_ = 1.50, p = 0.22, n=82; Figure 3B) did not affect tIgG. Plasma tIgG concentration did not differ among the different groups of adult females in Sonora and Jalisco (F_4,57_ = 0.60, p = 0.66, n=62; Figure 3C). In contrast, plasma tIgG concentration differed among age categories in Sonora (F_2,37_ = 13.26, p < 0.001, n=40; Figure 3D), with higher values in non-reproductive (p < 0.001) and lactating (p < 0.001) adult females than young.

**Fig 3.**
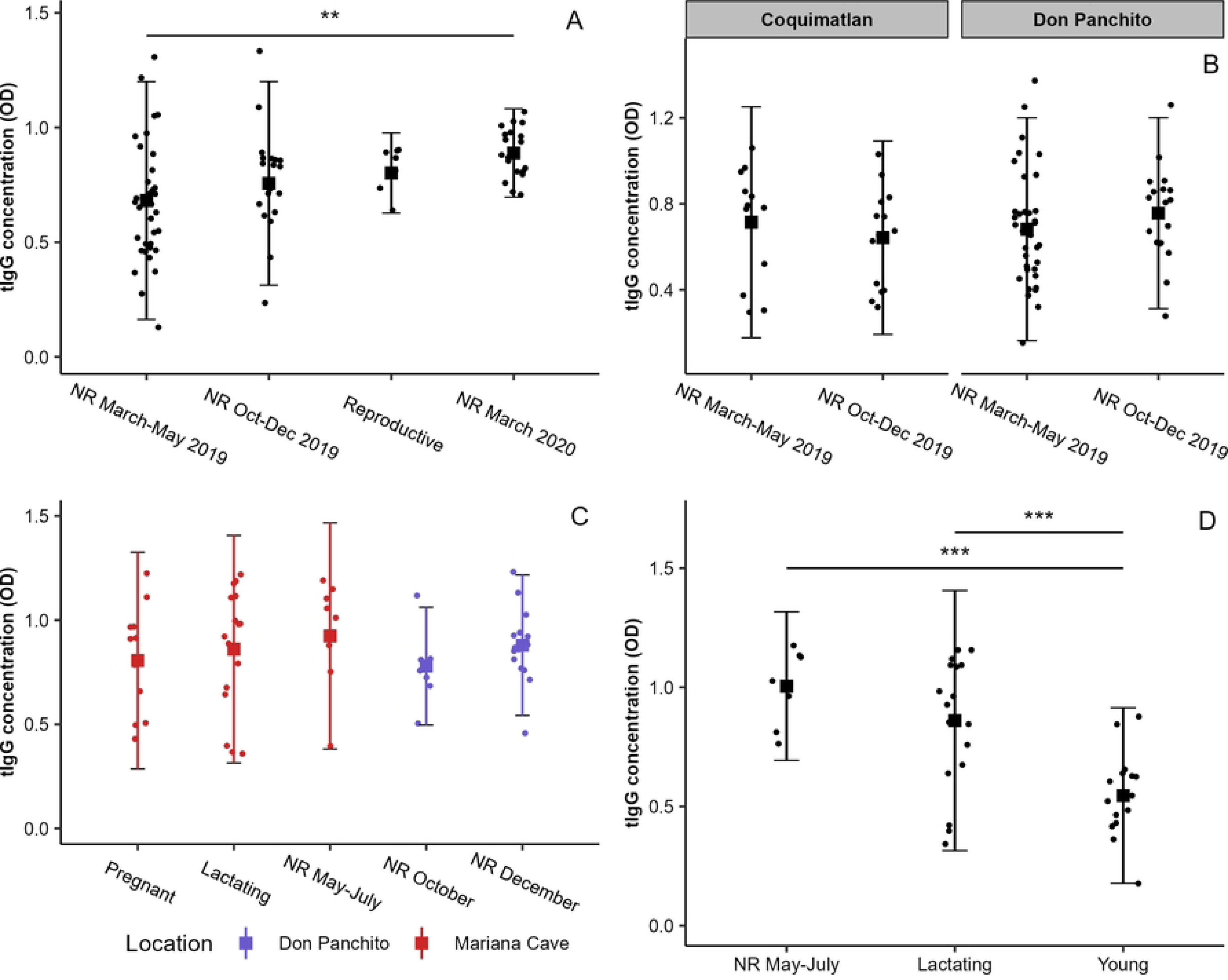
Total immunoglobulin G (tIgG) concentration in plasma across different demographic cohorts of the Lesser long-nosed bat (*Leptonycteris yerbabuenae*) in Mexico. A) reproductive and non-reproductive males in Chamela during March and May (NR March-May 2019) and October and December (NR Oct-Dec 2019) in 2019, and in March 2020 (NR March 2020), B) non-reproductive (NR) males of Coquimatlán and Chamela during the months of March and May (NR March-May) and October and December (NR Oct-Dec) in 2019, C) pregnant, lactating and non-reproductive females of Sonora (NR May-July), and non-reproductive females in October (NR October) and December (NR December) in Chamela, D) lactating, non-reproductive adult females (NR May-July) and young’s in Sonora.

### Correlations between plasma tIgG and BKA

The tIgG of non-reproductive males from Coquimatlán and Chamela did not correlate with the BKA (Spearman’s rho = -0.014, p = 0.561, n= 108; Figure 4A). In contrast, there was a positive correlation between these two measures in non-reproductive females (Spearman’s rho = 0.621, p < 0.001, n= 34; Figure 4B), and a nearly-significant correlation in reproductive females (Spearman’s rho = 0.279, p = 0.075, n= 28; Figure 4C).

**Fig 4.**
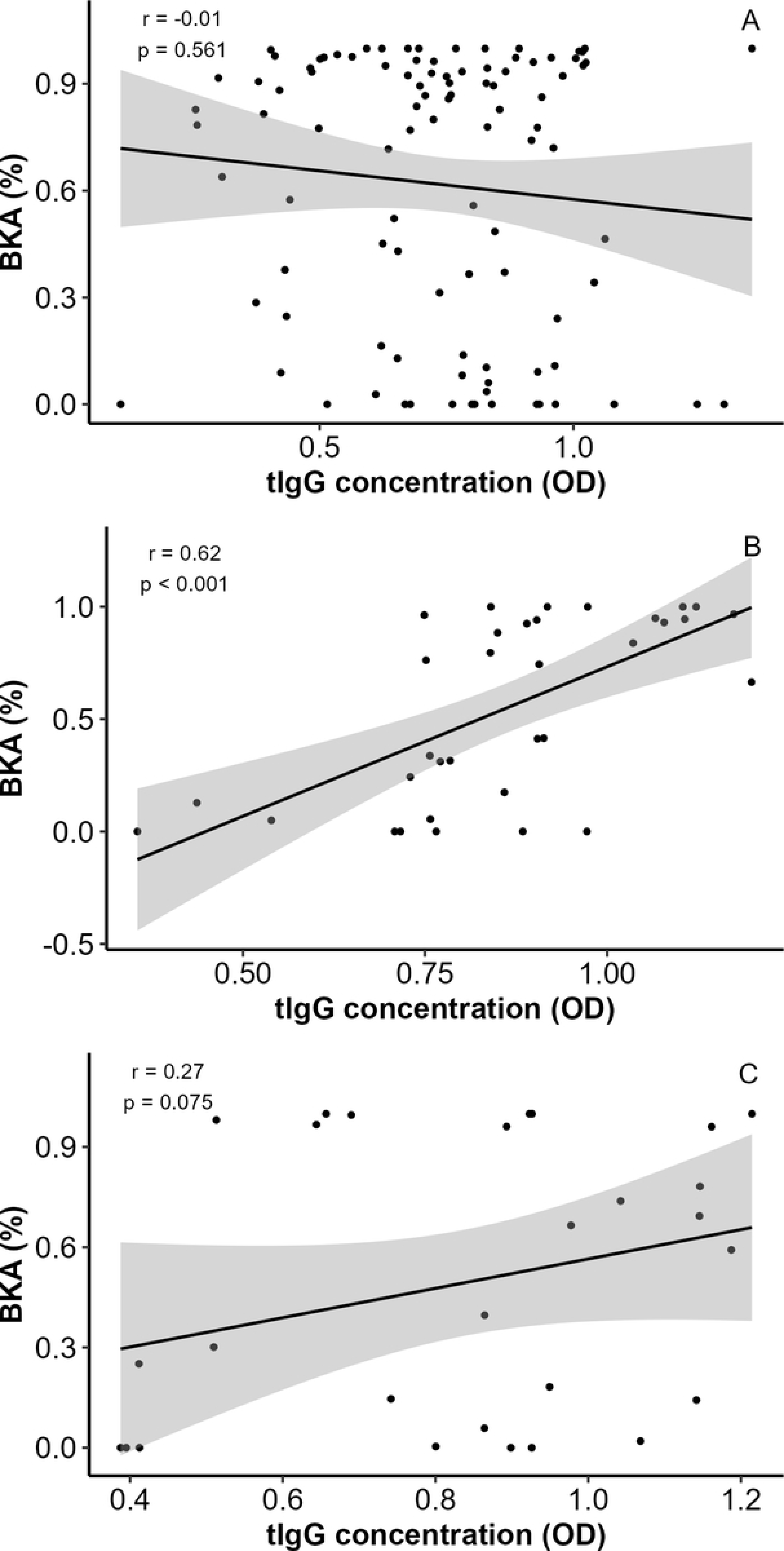
Plasma total IgG (tIgG) concentration and its relationship with bacterial-killing ability (BKA) of plasma in different demographic cohorts of the long-nosed bat (*L. yerbabuenae*) in Mexico. A) Non-reproductive males of Coquimatlán and Chamela, B) non-reproductive females and C) reproductive females.

## Discussion

Bats exhibit multiple life strategies to survive in different ecological contexts, which may influence the immune profile exhibited by different demographic cohorts. This relationship between immunity and demography in bats is controversial, understudied and key to identifying the possible factors that influence bat defenses and their relationship with the microorganisms they harbor. We provide evidence that tIgG concentration is stable throughout the adult reproductive cycle of *L. yerbabuenae* and is not affected by migration but is reduced in young individuals.

### Seasonal variation of tIgG concentration in adult males

In contrast to our prediction, reproductive males showed no difference in plasma tIgG concentrations compared to non-reproductive males. It has not been demonstrated that reproductive males of *L. yerbabuenae* experience an increase in testosterone concentration, so it is possible to postulate that reproductive season may not be directly related with an increase in testosterone, as is the case in some bat species [111]. However, we consider this scenario unlikely given the anatomical [80] and immunological [70] changes that *L. yerbabuenae* males undergo during the reproductive season. Another possible explanation is that antibody production is not affected by male reproduction in *L. yerbabuenae*, which would be consistent with the results obtained in *M. daubentonii* [44], but contradictory to the evidence observed in *D. rotundus* [45] and other vertebrate species in which testosterone diminishes antibody production [10,76,112–115]. Finally, it is possible that reproductively active males do not reflect the immunosuppressive effect of testosterone on antibody production, since the increased risk of infection during mating [116] and the bacterial load present in the dorsal patch of reproductive males [117] could stimulate tIgG production, because infection and antigenic stimulation are two of the main factors influencing plasma tIgG concentrations [77,118–123]. Finally, non-reproductive males had lower tIgG in March-May 2019 than in March 2020. There is no clear demographic factor that could explain these differences, suggesting that many individuals with reduced tIgG were sampled in March-May 2019. It is possible that these males with a reduced tIgG concentration may be experiencing an immunological trade-off if they were both infected and malnourished [124]. However, both cohorts did not exhibit differences in BMI, which suggest that this scenario is unlikely. Alternatively, tIgG concentration in some non-reproductive males in March-May 2019 could be related to changes in gut microbial composition, as gut microbiota is a key determinant of tIgG concentration [125].

### Effect of female reproduction and migration on tIgG concentration

The concentration of tIgG in females of *L. yerbabuenae* was stable throughout their reproductive cycle. Although the energetic cost of lactation is high [35,126] and may affect tIgG in some mammals [75,127], lactating females of *L. yerbabuenae* exhibited similar tIgG concentration than other female’s cohorts. This finding is consistent with the results reported for *M. daubentonii* [44] and with our hypothesis, because an energetic trade-off between lactation and tIgG is unlikely due to the high availability of resources in the area [64,81], the resilience of tIgG to caloric and nutritional deficits [128–130], and the fact that the effect of lactation on tIgG is limited to the first days post parturition [131–135]. Pregnancy in *L. yerbabuenae* did not affect tIgG, contradicting our hypothesis that tIgG concentration might increase as in *M. daubentonii* [44]. Other mammal species exhibit a decrease in maternal tIgG during the final stages of gestation [136–139] because the transport of systemic IgG to the fetus [140] or mammary glands [141]. However, llamas and alpacas did not exhibit changes in maternal tIgG around parturition [142], which is consistent with our findings because it is very likely that pregnant females sampled in this study were in late gestational stages [60,63]. To our knowledge, the type of placentation of *L. yerbabuenae* is unknown. These results suggest that it’s unlikely that *L. yerbabuenae* exhibits a type of hemochorial placentation, in which the transport of IgG to the fetus [140,143] reduces maternal tIgG [136,138]. Therefore, it is likely that IgG transfer occurs mainly via colostrum, in which case maternal tIgG remains constant throughout most of gestation and is only reduced at parturition [131,133,139]. However, we have no record of the hormonal profile of these females to accurately determine the stage of gestation and would need to sample females at parturition and take colostrum samples to confirm this hypothesis.

Migration can negatively affect tIgG in bats, as in the case of short-distance migrants of *L. noctivagans*, which may reflect an immunological trade-off with migration [58]. It is unlikely that the energy cost of migration compromises the tIgG in *L. yerbabuenae*, given the resilience of this antibody to energetic restraints [128–130] and the fact that this species is adapted to large daily movements [144]. We hypothesized that it would be possible to observe an increase in tIgG in migratory females during their return journey to west-central Mexico, as the use of multiple caves during this journey could increase the level of infection [145,146]. However, the results of this study suggest that tIgG is not affected at any stage of the migratory and reproductive cycle of *L. yerbabuenae* females, in contrast to what is observed with BKA [70]. To fully evaluate the effect of migration on tIgG in *L. yerbabuenae*, it is necessary to capture females that reside annually in the central-western region of Mexico [60,147], which was not considered in this study because we did not capture resident females.

Although the concentration of tIgG may reflect the presence of an infectious process [77,118–123], specific serological monitoring of potentially pathogenic microorganisms circulating within *L. yerbabuenae* colonies is essential to determine with certainty the effect of migration and reproduction on the risk of infection.

### Concentration of tIgG during early life

According to our hypothesis, lactating and non-reproductive adult females from the Sonoran region had a higher tIgG concentration than young, which is consistent with the positive relationship between age and tIgG observed in *S. bilineata* [96]. However, the minimum age (around 1 year) of *S. bilineata* used in Schneeberger et al 2014 [96] is likely not fully composed of young individuals [148,149]. Therefore, to our knowledge this is the first study to assess tIgG concentrations in bats during early-life, specifically in bats between 2 and 3 months of age because forearm length (53.17 mm ± 1.27) and weight (21.81g ± 1.64) were slightly higher than individuals of *L. yerbabuenae* with a recorded age of 35 days [150] and they were sampled 2-3 months after maternity colony formation in Sonora [62–64]. Consistent with this idea, young had a lower BMI than lactating and non-reproductive adult females in Sonora, suggesting that they are still in a growth phase.

The lower concentration of tIgG in young compared to adults is an immunological feature common to all mammals studied to date [71,73–75], regardless of the mode of placentation and maternal antibody transfer [143]. It is therefore plausible to argue that the reduced antibody production capacity of humans [31,151,152] and model rodents [153] during early life may also account for the reduced tIgG concentration in young bats of *L. yerbabuenae*. Consistent with this idea, the temporal dynamics of B lymphocytes in *R. aegyptiacus* bats resemble the changes in humans[32]. These ontogenetic changes in the immune system are closely linked to the immunological experience gained through exposure to environmental antigens [154]. Although some populations of B1 cells secrete natural IgG-type antibodies spontaneously and independently of antigenic stimuli [155], the main source of tIgG comes from plasma cells that have differentiated from antigen specific B2 lymphocytes [156–158]. Therefore, the reduced concentration of tIgG in young *L. yerbabuenae* most likely reflects the low degree of immunological experience they have compared to migratory adult females.

### Correlation between BKA and tIgG concentration in adult males and females

BKA is an immunological assay associated with innate immunity, which may have an antagonistic relationship with some parameters associated with acquired immunity in situations where the risk of infection or the energy status of the individual prioritizes the activation of one component over the other [45,159–161]. On the other hand, BKA is mainly dependent on complement activity [79,162–164], which can be activated by some IgG isotypes via the classical pathway [78,165,166]. In contrast to males, non-reproductive females showed a strong positive correlation between BKA and tIgG. These results suggest that non-reproductive females have a pool of IgG that recognizes *E. coli* and favors their elimination via the complement system. The presence of these antibodies in plasma could be a consequence of a bacterial translocation process from the gut to the blood [167,168], since *E. coli* inhabits the intestinal tract of bats [169,170] and its translocation to the blood could have stimulated the generation of antibodies [171]. However, it is also possible that antibodies directed against other types of microorganisms may cross-react with *E. coli* antigens [172,173], so it is essential to use single antigens against *E. coli* as a target to test this hypothesis. In addition to the existence of IgG that recognize *E. coli*, it is also plausible suggest that this relationship between BKA and tIgG in non-reproductive females could be a consequence of an IgG profile whose allotypes, isotypes and glycosylation promote complement activation [174,175].

## Conclusion

The results of this research suggest that plasma tIgG concentration is reduced during early life in *L. yerbabuenae*, and although systemic concentration of this antibody may be compromised in some adult individuals, tIgG levels are not affected by the reproductive season in adults and, in the case of females, by migration. Despite low tIgG concentration in young and the stability of tIgG levels in adults, there is a large variability between individuals, which may be a consequence of the genes [176] and their microbiota [125,167]. Therefore, future studies aimed at determining the contribution of these factors on tIgG in healthy individuals of *L. yerbabuenae* would allow the establishment of a reference range that would help to identify not only individuals who may be experiencing an infectious process [77,118–123], but also individuals who may be more resistant to infection [177,178]. In line with the latter idea, non-reproductive females showed a positive relationship between tIgG concentration and BKA, suggesting that females have a systemic antibody pool that enhances *E. coli* clearance. These results highlight the need for a thorough characterization of the IgG profile in *L. yerbabuenae*, as the subclasses present, their biological function and the antigens that stimulated their development are unknown.

## Supporting information

S1 Fig. Plasma bacteria-killing ability in non-reproductive males of the lesser long-nosed bats (*Leptonycteris yerbabuenae*).

S2 Fig. Plasma bacteria-killing ability of females of the lesser long-nosed bats (*Leptonycteris yerbabuenae*).

S3 Fig. Mean and standard deviation of IgG concentration in plasma samples of the lesser long-nosed bats (*Leptonycteris yerbabuenae*).

S1 Table. Average absorbance (490 nm), standard deviation and slope of total IgG concentration through two-fold serial dilutions of 10 randomly selected plasma samples of *Leptonycteris yerbabuenae*.

## Acknowledgments

The Ethics Committee in Research and Teaching of the Institute of Biology, National Autonomous University of Mexico (UNAM) approved the protocol of the study. The study was conducted under permit from Dirección General de Vida Silvestre (#8053/199). The study was supported by a research grant from Programa de Apoyo a Proyectos de Investigación e InnovaciónTecnológica, Dirección General de Asuntos del Personal Académico, UNAM (#IN203822) to L. Gerardo Herrera M. David A. Rivera-Ruiz was supported with a student grant from Secretaría de Ciencia, Humanidades, Tecnología e Innovación. We thank the logistic support of the personnel of the Chamela Biological Station in Jalisco, Natalia S. Herrera, León A. Pizano and Omar Calva for their help during field work, to Gastón Contreras Jiménez (Laboratorio de Microscopía y Microdisección Láser del Instituto de Ecología, UNAM) for his technical assistance in determining IgG concentration in the laboratory, and to José Ramírez M. for providing safe and enjoyable transportation to the island.

## References

1. wasaki A, Medzhitov R. Control of adaptive immunity by the innate immune system. Vol. 16, Nature Immunology. Nature Publishing Group; 2015. p. 343–53.

2. Zheng D, Liwinski T, Elinav E. Interaction between microbiota and immunity in health and disease. Cell Res [Internet]. 2020;30(6):492–506. Available from: 10.1038/s41422-020-0332-7

3. Pulendran B, Palucka K, Banchereau J. Sensing Pathogens and Tuning Immune Responses. Science (1979). 2001;293:253–6.

4. Maynard CL, Elson CO, Hatton RD, Weaver CT. Reciprocal interactions of the intestinal microbiota and immune system. Nature. 2012;489(7415):231–41.

5. Martin LB, Weil ZM, Nelson RJ. Seasonal changes in vertebrate immune activity : mediation by physiological trade-offs. Phil Trans R Soc B. 2008;363:321–39.

6. Adamo SA. The importance of physiology for ecoimmunology. In: Demas GE, Nelson RJ, editors. ECOIMMUNOLOGY. New York: Oxford University Press, Inc; 2012. p. 413–39.

7. Tieleman BI. Understanding immune function as a pace of life trait requires environmental context. Behav Ecol Sociobiol. 2018;72(55).

8. Demas GE, Greives T, Chester E, French S. The Energetics of Immunity. In: ECOIMMUNOLOGY. New York: Oxford University Press; 2012. p. 259–96.

9. Simon AK, Hollander GA, Mcmichael A. Evolution of the immune system in humans from infancy to old age. Proc R Soc B. 2015;282(20143085).

10. Klein SL, Flanagan KL. Sex differences in immune responses. Nat Rev Immunol. 2016;16:626–38.

11. Hegemann A, Fudickar AM, Nilsson JÅ. A physiological perspective on the ecology and evolution of partial migration. J Ornithol [Internet]. 2019;160(3):893–905. Available from: 10.1007/s10336-019-01648-9

12. Kunz TH, Fenton MB, editors. Bat Ecology. Chicago: University Chicago Press; 2003.

13. Schneeberger K, Czirják GÁ, Voigt CC. Measures of the Constitutive Immune System Are Linked to Diet and Roosting Habits of Neotropical Bats. PLoS One. 2013;8(1).

14. Allen LC, Turmelle AS, Mendonça MT, Navara KJ, Kunz TH, McCracken GF. Roosting ecology and variation in adaptive and innate immune system function in the Brazilian free-tailed bat (Tadarida brasiliensis). J Comp Physiol B. 2009;179:315–23.

15. Letko M, Seifert SN, Olival KJ, Plowright RK, Munster VJ. Bat-borne virus diversity, spillover and emergence. Vol. 18, Nature Reviews Microbiology. Nature Research; 2020. p. 461–71.

16. Szentivanyi T, McKee C, Jones G, Foster JT. Trends in Bacterial Pathogens of Bats: Global Distribution and Knowledge Gaps. Vol. 2023, Transboundary and Emerging Diseases. Wiley-Hindawi; 2023.

17. Roffler AA, Maurer DP, Lunn TJ, Sironen T, Forbes KM, Schmidt AG. Bat humoral immunity and its role in viral pathogenesis, transmission, and zoonosis. Front Immunol. 2024;15.

18. Banerjee A, Baker ML, Kulcsar K, Misra V, Plowright R, Mossman K. Novel Insights Into Immune Systems of Bats. Vol. 11, Frontiers in Immunology. Frontiers Media S.A.; 2020.

19. Plowright RK, Field HE, Smith C, Divljan A, Palmer C, Tabor G, et al. Reproduction and nutritional stress are risk factors for Hendra virus infection in little red flying foxes (Pteropus scapulatus). Proc R Soc B. 2008;275(1636):861–9.

20. Bergner LM, Orton RJ, Benavides JA, Becker DJ, Tello C, Biek R, et al. Demographic and environmental drivers of metagenomic viral diversity in vampire bats. Mol Ecol. 2020;29(1):26–39.

21. Dietrich M, Kearney T, Seamark ECJ, Paweska JT, Markotter W. Synchronized shift of oral, faecal and urinary microbiotas in bats and natural infection dynamics during seasonal reproduction. R Soc Open Sci. 2018;5(5).

22. Pereira M, Valério-Bolas A, Saraiva-Marques C, Alexandre-Pires G, da Fonseca IP, Santos-Gomes G. Development of dog immune system: From in uterus to elderly. Vet Sci. 2019;6(4).

23. Cardoso CL, King A, Chapwanya A, Esposito G. Ante-natal and post-natal influences on neonatal immunity, growth and puberty of calves—a review. Animals. 2021;11(5).

24. Adkins B, Leclerc C, Marshall-Clarke S. Neonatal adaptive immunity comes of age. Vol. 4, Nature Reviews Immunology. Nature Publishing Group; 2004. p. 553–64.

25. Georgountzou A, Nikolaos PG. Postnatal innate immune development : From birth to adulthood. Front Immunol. 2017;8(957):1–16.

26. Amman BR, Carroll SA, Reed ZD, Sealy TK, Balinandi S, Swanepoel R, et al. Seasonal Pulses of Marburg Virus Circulation in Juvenile Rousettus aegyptiacus Bats Coincide with Periods of Increased Risk of Human Infection. PLoS Pathog. 2012;8(10).

27. Storm N, Van Vuren PJ, Markotter W, Paweska JT. Antibody responses to marburg virus in egyptian rousette bats and their role in protection against infection. Viruses. 2018;10(2).

28. Epstein JH, Baker ML, Zambrana-Torrelio C, Middleton D, Barr JA, DuBovi E, et al. Duration of Maternal Antibodies against Canine Distemper Virus and Hendra Virus in Pteropid Bats. PLoS One. 2013;8(6):1–8.

29. Baker KS, Suu-Ire R, Barr J, Hayman DTS, Broder CC, Horton DL, et al. Viral antibody dynamics in a chiropteran host. Journal of Animal Ecology. 2014;83(2):415–28.

30. Sohayati AR, Hassan L, Sharifah SH, Lazarus K, Zaini CM, Epstein JH, et al. Evidence for Nipah virus recrudescence and serological patterns of captive Pteropus vampyrus. Epidemiol Infect. 2011;139(10):1570–9.

31. Labeur-Iurman L, Harker JA. Mechanisms of antibody mediated immunity – Distinct in early life. International Journal of Biochemistry and Cell Biology [Internet]. 2024;172(May):106588. Available from: 10.1016/j.biocel.2024.106588

32. Friedrichs V, Toussaint C, Schäfer A, Rissmann M, Dietrich O, Mettenleiter TC, et al. Landscape and age dynamics of immune cells in the Egyptian rousette bat. Cell Rep. 2022;40(10).

33. Martin SM, Malkinson TJ, Veale WL, Pittman QJ. Fever in pregnant, parturient, and lactating rats. American Journal of Physiology-Regulatory, Integrative and Comparative Physiology. 1995;268(64):R919–23.

34. Josefson CC, Zohdy S, Hood WR. Methodological Considerations for Assessing Immune Defense in Reproductive Females. Integr Comp Biol. 2020;60(3):732–41.

35. Speakman JR. The physiological costs of reproduction in small mammals. Phil Trans R Soc B. 2008;363:375–98.

36. Christe P, Arlettaz R, Vogel P. Variation in intensity of a parasitic mite (Spinturnix myoti) in relation to the reproductive cycle and immunocompetence of its bat host *(Myotis myotis)* . Ecol Lett. 2000;3:207–12.

37. Drexler JF, Corman VM, Wegner T, Tateno AF, Zerbinati RM, Gloza-Rausch F, et al. Amplification of Emerging Viruses in a Bat Bolony. Emerg Infect Dis. 2011;17(3):449–56.

38. Dietrich M, Wilkinson DA, Benlali A, Lagadec E, Ramasindrazana B, Dellagi K, et al. Leptospira and paramyxovirus infection dynamics in a bat maternity enlightens pathogen maintenance in wildlife. Environ Microbiol. 2015 Nov 1;17(11):4280–9.

39. Otálora-Ardila A, Flores-Martínez JJ, Rosales C, Salame-Méndez A, Herrera M. LG. Physiological and Ecological Correlates of the Cellular and Humoral Innate Immune Responses in an Insular Desert Bat: The Fish-Eating Myotis (Myotis vivesi). Diversity (Basel). 2022;14(781):1–17.

40. Li J, Chu Y, Yao W, Wu H, Feng J. Differences in Diet and Gut Microbiota Between Lactating and Non-lactating Asian Particolored Bats (Vespertilio sinensis): Implication for a Connection Between Diet and Gut Microbiota. Front Microbiol. 2021 Oct 12;12.

41. Phillips CD, Phelan G, Dowd SE, McDonough MM, Ferguson AW, Delton Hanson J, et al. Microbiome analysis among bats describes influences of host phylogeny, life history, physiology and geography. Mol Ecol. 2012 Jun;21(11):2617–27.

42. Gaona O, Gómez-Acata ES, Cerqueda-García D, Neri-Barrios CX, Falcón LI. Fecal microbiota of different reproductive stages of the central population of the lesser-long nosed bat, Leptonycteris yerbabuenae. PLoS One. 2019;14(7):1–19.

43. Hernández-Arciga U, Herrera M LG, Ibañez-Contreras A, Miranda-Labra RU, Flores-Martínez JJ, Königsberg M. Baseline and post-stress seasonal changes in immunocompetence and redox state maintenance in the fishing bat Myotis vivesi. PLoS One [Internet]. 2018;13(1):e0190047. Available from: 10.1371/journal.pone.0190047

44. Ruoss S, Becker NI, Otto MS, Czirják G, Encarnação JA. Effect of sex and reproductive status on the immunity of the temperate bat Myotis daubentonii. Mammalian Biology [Internet]. 2019;94:120–6. Available from: 10.1016/j.mambio.2018.05.010

45. Becker DJ, Czirják G, Volokhov D V., Bentz AB, Carrera JE, Camus MS, et al. Livestock abundance predicts vampire bat demography, immune profiles and bacterial infection risk. Phil Trans R Soc B. 2018;373(20170089).

46. Herrera LG, Hernández MU, Antonio J, Carcacía G, Nassar JM. Olfactory cues of mate quality in mammals : inflammatory response is higher in males of long - nosed bats with odorous dorsal patch. 2023;(April).

47. Gnanadesikan GE, Pearse WD, Shaw AK. Evolution of mammalian migrations for refuge, breeding, and food. Ecol Evol. 2017 Aug 1;7(15):5891–900.

48. Fudickar AM, Jahn AE, Ketterson ED. Animal Migration: An Overview of One of Nature’s Great Spectacles. The Annual Review of Ecology, Evolution, and Systematics [Internet]. 2021;52:17. Available from: 10.1146/annurev-ecolsys-012021-

49. Poulin R, de Angeli Dutra D. Animal migrations and parasitism: reciprocal effects within a unified framework. Biological Reviews. 2021;96(4):1331–48.

50. de Angeli Dutra D, Fecchio A, Martins Braga É, Poulin R. Migratory birds have higher prevalence and richness of avian haemosporidian parasites than residents. Int J Parasitol [Internet]. 2021;51(10):877–82. Available from: 10.1016/j.ijpara.2021.03.001

51. Hall RJ, Altizer S, Peacock SJ, Shaw AK. Animal migration and infection dynamics: Recent advances and future frontiers. In: Ezenwa V, Altizer SM, Hall R, editors. Animal Behavior and Parasitism [Internet]. online edn. Oxford University Press; 2022. p. 111-132; Available from: 10.1093/oso/9780192895561.003.0007

52. Altizer S, Bartel R, Han BA. Animal migration and infectious disease risk. Science (1979). 2011;331(6015):296–302.

53. Shaw AK, Binning SA. Recovery from infection is more likely to favour the evolution of migration than social escape from infection. Journal of Animal Ecology. 2020;89(6):1448–57.

54. Shaw AK, Binning SA. Migratory recovery from infection as a selective pressure for the evolution of migration. American Naturalist. 2016;187(4):502–16.

55. Owen JC, Moore FR. Relationship between energetic condition and indicators of immune function in thrushes during spring migration. Can J Zool. 2008;86(7):638– 47.

56. Eikenaar C, Isaksson C, Hegemann A. A hidden cost of migration? Innate immune function versus antioxidant defense. Ecol Evol. 2018;(November 2017):2721–8.

57. Voigt CC, Fritze M, Lindecke O, Costantini D, Pētersons G, Czirják G. The immune response of bats differs between pre-migration and migration seasons. Sci Rep [Internet]. 2020;10(17384). Available from: 10.1038/s41598-020-74473-3

58. Rogers EJ, McGuire L, Longstaffe FJ, Clerc J, Kunkel E, Fraser E. Relating wing morphology and immune function to patterns of partial and differential bat migration using stable isotopes. Journal of Animal Ecology. 2022 Apr 1;91(4):858–69.

59. Medellín R. Leptonycteris yerbabuenae. The IUCN red list of threatened species. [Internet]. The IUCN red list of threatened species. 2016 [cited 2020 Sep 9]. Available from: https://iucnredlist.org/species/136659/21988965

60. Stoner KE, Karla KA, Roxana RC, Quesada M. Population dynamics, reproduction, and diet of the lesser long-nosed bat (Leptonycteris curasoae) in Jalisco, Mexico: Implications for conservation. Biodivers Conserv. 2003;12(2):357–73.

61. Ceballos G, Fleming TH, Chavez C, Nassar J. Population Dynamics of Leptonycteris curasoae (Chiroptera: Phyllostomidae) in Jalisco, Mexico. J Mammal. 1997;78(4):1220–30.

62. Cockrum EL. Seasonal Distribution of Northwestern Populations of the Long-Nosed Bats, Leptonycteris sanborni Family Phyllostomidae. Anales del Instituto de Biología Serie Zoología [Internet]. 1991;62(2):181–202. Available from: https://www.redalyc.org/articulo.oa?id=45862206

63. Peñalba MC, Molina-Freaner F, Rodríguez LL. Resource availability, population dynamics and diet of the nectar-feeding bat Leptonycteris curasoae in Guaymas, Sonora, Mexico. Biodivers Conserv. 2006 Aug;15(9):3017–34.

64. Horner MA, Fleming TH, Sahley CT. Foraging behaviour and energetics of a nectar-feeding bat, Leptonycteris curasoae (Chiroptera: Phyllostomidae). J Zool [Internet]. 1998;244(4):575–86. Available from: doi:10.1111/j.1469-7998.1998.tb00062.x

65. Menchaca A, Arteaga MC, Medellin RA, Jones G. Conservation units and historical matrilineal structure in the tequila bat (Leptonycteris yerbabuenae). Glob Ecol Conserv. 2020;23.

66. Trejo-Salazar RE, Castellanos-Morales G, Hernández-Rosales DC, Gámez N, Gasca-Pineda J, Morales Garza MR, et al. Discordance in maternal and paternal genetic markers in lesser long-nosed bat leptonycteris yerbabuenae, a migratory bat: Recent expansion to the north and male phylopatry. PeerJ. 2021;9.

67. Wilkinson GS, Fleming TH. Migration and evolution of lesser longnosed bats Lepfonycteris curasoae, inferred from mitochondrial DNA. Mol Ecol. 1996;5:329– 39.

68. Arteaga MC, Medellín RA, Luna-Ortíz PA, Heady PA, Frick WF. Genetic diversity distribution among seasonal colonies of a nectar-feeding bat (Leptonycteris yerbabuenae) in the Baja California Peninsula. Mammalian Biology. 2018 Sep 1;92:78–85.

69. Morales-Garza MR, Arizmendi M del C, Campos JE, Martínez-Garcia M, Valiente-Banuet A. Evidences on the migratory movements of the nectar-feeding bat Leptonycteris curasoae in Mexico using random amplified polymorphic DNA (RAPD). J Arid Environ. 2007;68(2):248–59.

70. Rivera-Ruiz DA, Flores-Martínez JJ, Rosales C, Herrera Montalvo LG. Constitutive Innate Immunity of Migrant and Resident Long-Nosed Bats (Leptonycteris yerbabuenae) in the Drylands of Mexico. Diversity (Basel). 2023 Apr 1;15(4).

71. Bayram RO, Özdemir H, Emsen A, Türk Daği H, Artaç H. Reference ranges for serum immunoglobulin (IgG, IgA, and IgM) and IgG subclass levels in healthy children. Turki J Med Sci. 2019;49:497–505.

72. Jazayeri MH, Pourfathollah AA, Rasaee MJ, Porpak Z, Jafari ME. The concentration of total serum IgG and IgM in sera of healthy individuals varies at different age intervals. Biomedicine and Aging Pathology [Internet]. 2013;3(4):241–5. Available from: 10.1016/j.biomag.2013.09.002

73. Erhard MH, Amon P, Younan M, Ali Z, Stangassinger M. Absorption and synthesis of immunoglobulins G in newborn calves. Reproduction in Domestic Animals. 1999;34(3–4):173–5.

74. Chastant S, Mila H. Passive immune transfer in puppies. Anim Reprod Sci [Internet]. 2019;207(March):162–70. Available from: 10.1016/j.anireprosci.2019.06.012

75. King DP, Lowe K a, Hay AWM, Evans SW. Identification, characterisation, and measurement of immunoglobulin concentrations in grey (Haliocherus grypus) and common (Phoca vitulina) seals. Dev Comp Immunol. 1994;18(5):433–42.

76. Kanda N, Tsuchida T, Tamaki K. Testosterone inhibits immunoglobulin production by human peripheral blood mononuclear cells. Clin Exp Immunol. 1996;106(2):410– 5.

77. Abolins S, King EC, Lazarou L, Weldon L, Hughes L, Drescher P, et al. The comparative immunology of wild and laboratory mice, Mus musculus domesticus. Nat Commun [Internet]. 2017;8(14811):1–13. Available from: 10.1038/ncomms14811

78. Goldberg BS, Ackerman ME. Antibody-mediated complement activation in pathology and protection. Vol. 98, Immunology and Cell Biology. John Wiley and Sons Inc.; 2020. p. 305–17.

79. Moore MS, Reichard JD, Murtha TD, Zahedi B, Fallier RM, Thomas H. Specific Alterations in Complement Protein Activity of Little Brown Myotis (Myotis lucifugus) Hibernating in White-Nose Syndrome Affected Sites. PLoS One. 2011;6(11):e27430.

80. Rincón-Vargas F, Stoner KE, Vigueras-Villaseñor RM, Nassar JM, Chaves ÓM, Hudson R. Internal and external indicators of male reproduction in the lesser long-nosed bat Leptonycteris yerbabuenae. J Mammal. 2013;94(2):488–96.

81. Burke RA, Frey JK, Ganguli A, Stoner KE. Species distribution modelling supports “nectar corridor” hypothesis for migratory nectarivorous bats and conservation of tropical dry forest. Divers Distrib. 2019;25(9):1399–415.

82. Fleming TH, Nuñez RA, Sternberg L da SL. Seasonal changes in the diets of migrant and non-migrant nectarivorous bats as revealed by carbon stable isotope analysis. Oecologia. 1993;94(1):72–5.

83. Sikes RS, Gannon william L, Mammalogists the AC and UC of the AS of, L W. Guidelines of the American Society of Mammalogists for the use of wild mammals in research. J Mammal. 2011;92(1):235–53.

84. Anthony ELP. Age determination in bats. In: Kunz TH, editor. Ecological and Behavioral Methods for the Study of Bats. Washington, DC/ London: Smithsonian Institution Press: Washington; 1988. p. 47–58.

85. Brunet-Rossinni AK, Wilkinson GS. Methods for age estimation and the study of senescence in bats. In: Kunz TH, Parsons S, editors. Ecological and Behavioral Methods for the Study of Bats. 2nd ed. Baltimore, Maryland: Johns Hopkins University Press; 2009. p. 315–25.

86. Nassar JM, Salazar MV, Quintero A, Stoner KE, Gómez M, Cabrera A, et al. Seasonal sebaceous patch in the nectar-feeding bats Leptonycteris curasoae and L. yerbabuenae (Phyllostomidae: Glossophaginae): phenological, histological, and preliminary chemical characterization. Zoology. 2008;111(5):363–76.

87. Muñoz-Romo M, Nielsen LT, Nassar JM, Kunz TH. Chemical composition of the substances from dorsal patches of males of the Curaçaoan long-nosed bat, Leptonycteris curasoae (Phyllostomidae: Glossophaginae). Acta Chiropt. 2012;14(1):213–24.

88. Park KJ, Jones G, Ransome RD. Torpor, Arousal and Activity of Hibernating Greater Horseshoe Bats (Rhinolophus ferrumequinum). Ecology [Internet]. 2000 [cited 2024 May 17];14(5):580–8. Available from: http://www.jstor.org/stable/2656391

89. Pearce RD, O’Shea TJ, Wunder BA. Evaluation of morphological indices and total body electrical conductivity to assess body composition in big brown bats. Acta Chiropt. 2008;10(1):153–9.

90. Speakman JR, Racey PA. The influence of body condition on sexual development of male Brown long-eared bats (Plecotus auritus) in the wild. J Zool. 1986;210(4):515– 25.

91. Lacki MJ, Dodd LE, Toomey RS, Thomas SC, Couch ZL, Nichols BS. Temporal changes in body mass and body condition of cave-hibernating bats during staging and swarming. J Fish Wildl Manag. 2015 Dec 1;6(2):360–70.

92. Durán AA. The body mass index is influenced by the environment in populations of Artibeus planirostris (Chiroptera: Phyllostomidae)? Acta Zoologica Lilloana. 2022 Jun 1;66(1):1–9.

93. McGuire LP, Kelly LA, Baloun DE, Boyle WA, Cheng TL, Clerc J, et al. Common condition indices are no more effective than body mass for estimating fat stores in insectivorous bats. J Mammal. 2018 Oct 10;99(5):1065–71.

94. Liebl AL, Ii LBM. Simple quantification of blood and plasma antimicrobial capacity using spectrophotometry. Funct Ecol. 2009;23:1091–6.

95. French SS, Neuman-lee LA. Improved ex vivo method for microbiocidal activity across vertebrate species. Biol Open. 2012;

96. Schneeberger K, Courtiol A, Czirja Gabor A, Voigt CC. Immune Profile Predicts Survival and Reflects Senescence in a Small, Long-Lived Mammal, the Greater Sac-Winged Bat (Saccopteryx bilineata). PLoS One. 2014;9(9).

97. Strobel SN, Becker JA. No short-term effect of handling and capture stress on immune responses on bats assessed by bacterial killing assay. Mammalian Biology [Internet]. 2015; Available from: 10.1016/j.mambio.2015.02.005

98. Becker DJ, Chumchal MM, Bentz AB, Platt SG, Czirják G, Rainwater TR, et al. Predictors and immunological correlates of sublethal mercury exposure in vampire bats. R Soc Open Sci. 2017;4(170073).

99. Seltmann A, Troxell SA, Schad J, Fritze M, Bailey LD, Voigt CC, et al. Differences in acute phase response to bacterial, fungal and viral antigens in greater mouse-eared bats (Myotis myotis). Sci Rep. 2022 Dec 1;12(1).

100. Fritze M, Puechmaille SJ, Costantini D, Fickel J, Voigt CC, Czirják G. Determinants of defence strategies of a hibernating European bat species towards the fungal pathogen Pseudogymnoascus destructans. Dev Comp Immunol. 2021 Jun 1;119.

101. Costantini D, Czirják G, Bustamante P, Bumrungsri S, Voigt CC. Impacts of land use on an insectivorous tropical bat: The importance of mercury, physio-immunology and trophic position. Science of the Total Environment. 2019 Jun 25;671:1077–85.

102. Ingala MR, Becker DJ, Bak Holm J, Kristiansen K, Simmons NB. Habitat fragmentation is associated with dietary shifts and microbiota variability in common vampire bats. Ecol Evol. 2019;9(11):6508–23.

103. R Core Team. R: A language and environment for statistical computing, Vienna, Austria. Vienna, Austria: R Foundation for Statistical Computing; 2022.

104. Delignette-Muller ML, Dutang C. fitdistrplus: An R Package for Fitting Distributions. J Stat Softw. 2015;64(4):1–34.

105. Fox J, Weisberg S. An R Companion to Applied Regression [Internet]. Third. Thousand Oaks, Canada: Sage; 2019. Available from: https://socialsciences.mcmaster.ca/jfox/Books/Companion/%0A

106. Hothorn T, Bretz F, Westfall P. Simultaneous Inference in General Parametric Models. Biometrical Journal. 2008;50(3):346–63.

107. Lenth R. emmeans: Estimated marginal means, aka least-squares means [Internet]. R package version 1.8.9; 2023 [cited 2024 Nov 5]. Available from: https://CRAN.R-project.org/package=emmeans

108. Miller GA, Chapman JP. Misunderstanding analysis of covariance. J Abnorm Psychol. 2001;110(1):40–8.

109. Hartig F. DHARMa: Residual Diagnostics for Hierarchical (Multi-Level/Mixed) regression models [Internet]. R package version 0.4.6; 2022. Available from: https://cran.r-project.org/package=DHARMa

110. Puth MT, Neuhäuser M, Ruxton GD. Effective use of Spearman’s and Kendall’s correlation coefficients for association between two measured traits. Vol. 102, Animal Behaviour. Academic Press; 2015. p. 77–84.

111. Martin L, Bernard RTF. Endocrine Regulation of Reproduction in Bats: The Role of Circulating Gonadal Hormones. In: Crichton EG, Krutzsch PH, editors. Reproductive Biology of Bats [Internet]. Academic Press; 2000. p. 27–64. Available from: http://www.sciencedirect.com/science/article/pii/B9780121956707500035

112. Derting TL, Virk ÆMK. Positive effects of testosterone and immunochallenge on energy allocation to reproductive organs. J Comp Physiol B. 2005;543–56.

113. Greives TJ, McGlothlin JW, Jawor JM, Demas GE, Ketterson ED. Testosterone and Innate Immune Function Inversely Covary in a Wild Population of Breeding Dark-Eyed Juncos (Junco hyemalis). Funct Ecol. 2006;20(5):812–8.

114. Daniels CW, Belosevic M. Serum antibody responses by male and female C57B1/6 mice infected with Giardia muris. Clin Exp Immunol. 1994;97:424–9.

115. Kanda N, Tsuchida T, Tamaki K. Testosterone suppresses anti-DNA antibody production in peripheral blood mononuclear cells from patients with systemic lupus erythematosus. Arthritis Rheum. 1997 Sep;40(9):1703–11.

116. Nunn CL, Gittleman JL, Antonovics J. Promiscuity and the primate immune system. Science (1979). 2000;290(5494):1168–70.

117. Gaona O, Cerqueda-García D, Falcón LI, Vázquez-Domínguez G, Valdespino-Castillo PM, Neri-Barrios CX. Microbiota composition of the dorsal patch of reproductive male Leptonycteris yerbabuenae. PLoS One. 2019;14(12):1–15.

118. Abolins SR, Pocock MJO, Hafalla JCR, Riley EM, Viney ME. Measures of immune function of wild mice, Mus musculus. Mol Ecol. 2011;20(5):881–92.

119. Maden M, Birdane FM, Uçan US, Altunok V. Concentrations of total serum immunoglobulin E, A, G and M in stray dogs with healthy and dermatological problems. Kafkas Univ Vet Fak Derg. 2013;19(2):347–50.

120. Proverbio D, Spada E, Bagnagatti De Giorgi G, Perego R, Valena E. Relationship between Leishmania IFAT titer and clinicopathological manifestations (clinical score) in dogs. Biomed Res Int. 2014;2014.

121. Hunziker L, Recher M, Macpherson AJ, Ciurea A, Freigang S, Hengartner H, et al. Hypergammaglobulinemia and autoantibody induction mechanisms in viral infections. Nat Immunol. 2003 Apr 1;4(4):343–9.

122. Cave NJ, Bridges JP, Thomas DG. Systemic effects of periodontal disease in cats. Veterinary Quarterly. 2012 Sep;32(3–4):131–44.

123. Beuvon C, Martin M, Baillou C, Roblot P, Puyade M. Etiologies of Polyclonal Hypergammaglobulinemia: A scoping review. Vol. 90, European Journal of Internal Medicine. Elsevier B.V.; 2021. p. 119–21.

124. Michael H, Langel SN, Miyazaki A, Paim FC, Chepngeno J, Alhamo MA, et al. Malnutrition Decreases Antibody Secreting Cell Numbers Induced by an Oral Attenuated Human Rotavirus Vaccine in a Human Infant Fecal Microbiota Transplanted Gnotobiotic Pig Model. Front Immunol. 2020 Feb 14;11.

125. Kim M, Qie Y, Park J, Kim CH. Gut Microbial Metabolites Fuel Host Antibody Responses. Cell Host Microbe. 2016 Aug 10;20(2):202–14.

126. Dewey KG. Energy and protein requirements during lactation. Annu Rev Nutr [Internet]. 1997;17:19–36. Available from: www.annualreviews.org

127. Flies AS, Mansfield LS, Flies EJ, Grant CK, Holekamp KE. Socioecological predictors of immune defences in wild spotted hyenas. Funct Ecol. 2016;30(9):1549–57.

128. Rytter MJH, Kolte L, Briend A, Friis H, Christensen VB. The immune system in children with malnutrition - A systematic review. PLoS One. 2014 Aug 25;9(8).

129. Brüssow H, Barclay D, Sidoti J, Rey S, Blondel A, Dirren H, et al. Effect of Malnutrition on Serum and Milk Antibodies in Zairian Women. Clin Diagn Lab Immunol [Internet]. 1996;3(1):37–41. Available from: https://journals.asm.org/journal/cdli

130. Brown RF, Bartrop R, Birmingham CL. Immunological disturbance and infectious disease in anorexia nervosa: A review. Vol. 20, Acta Neuropsychiatrica. Blackwell Publishing Ltd; 2008. p. 117–28.

131. Hernández-Castellano LE, Moreno-Indias I, Sánchez-Macías D, Morales-delaNuez A, Torres A, Argüello A, et al. Sheep and goats raised in mixed flocks have diverse immune status around parturition. J Dairy Sci [Internet]. 2019;102(9):8478–85. Available from: 10.3168/jds.2019-16731

132. Herr M, Bostedt H, Failing K. IgG and IgM levels in dairy cows during the periparturient period. Theriogenology [Internet]. 2011;75(2):377–85. Available from: 10.1016/j.theriogenology.2010.09.009

133. Anugu S, Petersson-Wolfe CS, Combs GF, Petersson KH. Effect of vitamin E on the immune system of ewes during late pregnancy and lactation. Small Ruminant Research. 2013 Apr;111(1–3):83–9.

134. Devillers N, Farmer C, Mounier AM, Le Dividich J, Prunier A. Hormones, IgG and lactose changes around parturition in plasma, and colostrum or saliva of multiparous sows. Reprod Nutr Dev. 2004 Jul;44(4):381–96.

135. Czyżewska-Dors E, Wierzchosławski K, Pomorska-Mól M. Serum concentrations of immunoglobulins and cortisol around parturition in clinically healthy sows and sows with postpartum dysgalactia syndrome (PDS). J vet Res. 2022 Jun 1;66(2):245–50.

136. Zhang T, Hu Y, Xiang Z. Changes of serum immunoglobulin level in healthy pregnant women and establishment of its reference interval. Journal of Central South University (Medical Sciences). 2021 Jan 1;46(1):53–9.

137. Castro N, Capote J, Martín D, Argüello A. The influence of dietary conjugated linoleic acid on blood serum and colostrum immunoglobulin G concentration in female goats before and after parturition. J Anim Physiol Anim Nutr (Berl). 2006;90(9–10):429–31.

138. Gustafsson E, Mattsson A, Holmdahl R, Mattsson R. Pregnancy in B-cell-deficient mice: postpartum transfer of immunoglobulins prevents neonatal runting and death. Biol Reprod [Internet]. 1994;51(6):1173–80. Available from: https://academic.oup.com/biolreprod/article/51/6/1173/2761421

139. Klobasa F, Habe F, Werhahn E, Butler JE. Changes in the concentrations of serum IgG, IgA and IgM of sows throughout the reproductive cycle. Vet Immunol Immunopathol. 1985;10(4):341–53.

140. Palmeira P, Quinello C, Silveira-Lessa AL, Zago CA, Carneiro-Sampaio M. IgG placental transfer in healthy and pathological pregnancies. Vol. 2012, Clinical and Developmental Immunology. 2012.

141. Hine BC, Hunt PW, Colditz IG. Production and active transport of immunoglobulins within the ruminant mammary gland. Vol. 211, Veterinary Immunology and Immunopathology. Elsevier B.V.; 2019. p. 75–84.

142. Bravo PW, Garnica J, Fowler ’ ME. Immunoglobulin G concentrations in periparturient llamas, alpacas and their crias. Small Ruminant Research. 1997;26:145–9.

143. Borghesi J, Mario LC, Rodrigues MN, Favaron PO, Miglino MA. Immunoglobulin Transport during Gestation in Domestic Animals and Humans—A Review. Open J Anim Sci. 2014;04(05):323–36.

144. Medellin RA, Rivero M, Ibarra A, De La Torre JA, Gonzalez-Terrazas TP, Torres-Knoop L, et al. Follow me: Foraging distances of Leptonycteris yerbabuenae (Chiroptera: Phyllostomidae) in Sonora determined by fluorescent powder. J Mammal. 2018;99(2):306–11.

145. Li L, Victoria JG, Wang C, Jones M, Fellers GM, Kunz TH, et al. Bat Guano Virome: Predominance of Dietary Viruses from Insects and Plants plus Novel Mammalian Viruses. J Virol [Internet]. 2010;84(14):6955–65. Available from: http://jvi.asm.org/cgi/doi/10.1128/JVI.00501-10

146. Banskar S, Bhute SS, Suryavanshi M V., Punekar S, Shouche YS. Microbiome analysis reveals the abundance of bacterial pathogens in Rousettus leschenaultii guano. Sci Rep. 2016;6(36948):1–13.

147. Rojas-Martinez A, Valiente-Banuet A, Arizmendi M del coro, Alcántara-Eguren A, Arita HT. Seasonal distribution of the long-nosed bat (Leptonycteris curasoae) in North America: does a generalized migration pattern really exist? J Biogeogr. 1999;26:1065–77.

148. Yancey FD, Goetze JR, Jones C. Saccopteryx bilineata. Mammalian Species. 1998;(581):1–5.

149. Strauss M, Helversen O Von, Knörnschild M. The ontogeny of courtship behaviours in bat pups (Saccopteryx bilineata). Behaviour. 2010;147(5):661–76.

150. Martínez-Coronel M, Cervantes FA, Hortelano-Moncada Y. Crecimiento postnatal y desarrollo del vuelo en el murciélago Leptonycteris yerbabuenae en Chiapas, México. Therya. 2014;5(1):303–22.

151. Clemens EA, Alexander-Miller MA. Understanding antibody responses in early life: Baby steps towards developing an effective influenza vaccine. Viruses. 2021;13(7).

152. Hill DL, Carr EJ, Rutishauser T, Moncunill G, Campo JJ, Innocentin S, et al. Immune system development varies according to age, location, and anemia in African children. Sci Transl Med. 2020;12(529).

153. Pyle CJ, Labeur-Iurman L, Groves HT, Puttur F, Lloyd CM, Tregoning JS, et al. Enhanced IL-2 in early life limits the development of Tfh and protective antiviral immunity. Journal of Experimental Medicine. 2021;218(12).

154. Pia̧tosa B, Wolska-Kuśnierz B, Pac M, Siewiera K, Gałkowska E, Bernatowska E. B cell subsets in healthy children: Reference values for evaluation of B cell maturation process in peripheral blood. Cytometry B Clin Cytom. 2010 Nov;78(6 B):372–81.

155. Palma J, Tokarz-Deptuła B, Deptuła J, Deptuła W. Natural antibodies – Facts known and unknown. Central European Journal of Immunology. 2018;43(4):466–75.

156. Manz RA, Hauser AE, Hiepe F, Radbruch A. Maintenance of serum antibody levels. Vol. 23, Annual Review of Immunology. 2005. p. 367–86.

157. Bos NA, Kimura H, Meeuwsen CG, Visser H De, Hazenberg MP, Wostmann BS, et al. Serum immunoglobulin levels and naturally occurring antibodies against carbohydrate antigens in germ-free BALB/c mice fed chemically defined ultrafiltered diet. Eur J Immunol. 1989;19(12):2335–9.

158. Cukrowska B, Kozáková H, Eháková ZŘ, Inkora JŠ, Tlaskalová-Hogenová H.Specific Antibody and Immunoglobulin Responses after Intestinal Colonization of Germ-Free Piglets with Non-Pathogenic Escherichia coli O86. Immunobiology [Internet]. 2001;204:425–33. Available from: http://www.urbanfischer.de/journals/immunobiol

159. Lee KA. Linking immune defenses and life history at the levels of the individual and the species. Integr Comp Biol. 2006;46(6):1000–15.

160. Mcdade TW, Georgiev A V, Kuzawa CW. Trade-offs between acquired and innate immune defenses in humans. Evol Med Public Health. 2016;1–16.

161. Ezenwa VO, Ekernas LS, Creel S. Unravelling complex associations between testosterone and parasite infection in the wild. Funct Ecol. 2012;26:123–33.

162. Merchant ME, Roche C, Elsey RM, Prudhomme J. Antibacterial properties of serum from the American alligator (Alligator mississippiensis). Comparative Biochemistry and Physiology Part B. 2003;136:505–13.

163. Matson KD, Ricklefs RE, Klasing KC. A hemolysis-hemagglutination assay for characterizing constitutive innate humoral immunity in wild and domestic birds. Dev Comp Immunol. 2005;29(3):275–86.

164. Becker DJ, Czirják GÁ, Rynda-apple A, Plowright RK. Handling Stress and Sample Storage Are Associated with Weaker Complement-Mediated Bactericidal Ability in Birds but Not Bats. Physiological and Biochemical Zoology. 2019;92(1):37–48.

165. Merle NS, Church SE, Fremeaux-Bacchi V, Roumenina LT. Complement system part I - molecular mechanisms of activation and regulation. Vol. 6, Frontiers in Immunology. Frontiers Research Foundation; 2015.

166. Nesargikar P, Spiller B, Chavez R. The complement system: History, pathways, cascade and inhibitors. Eur J Microbiol Immunol (Bp). 2012 Jun;2(2):103–11.

167. Vujkovic-Cvijin I, Welles HC, Ha CWY, Huq L, Mistry S, Brenchley JM, et al. The systemic anti-microbiota IgG repertoire can identify gut bacteria that translocate across gut barrier surfaces [Internet]. Vol. 14, Sci. Transl. Med. 2022. Available from: https://www.science.org

168. Zeng MY, Cisalpino D, Varadarajan S, Hellman J, Warren HS, Cascalho M, et al. Gut Microbiota-Induced Immunoglobulin G Controls Systemic Infection by Symbiotic Bacteria and Pathogens. Immunity. 2016;44(3):647–58.

169. Nowakiewicz A, Zięba P, Gnat S, Trościańczyk A, Osińska M, Dominik Ł, et al. Bats as a reservoir of resistant Escherichia coli: A methodical view. Can we fully estimate the scale of resistance in the reservoirs of free-living animals? Res Vet Sci. 2020;128:49–58.

170. Federici L, Masulli M, De Laurenzi V, Allocati N. An overview of bats microbiota and its implication in transmissible diseases. Vol. 13, Frontiers in Microbiology. Frontiers Media S.A.; 2022.

171. Liu X, Peng X, Li H. Escherichia coli Activate Extraintestinal Antibody Response and Provide Anti-Infective Immunity. Int J Mol Sci. 2024 Jul 1;25(13).

172. Augustyniak D, Majkowska-Skrobek G, Roszkowiak J, Dorotkiewicz-Jach A. Defensive and Offensive Cross-Reactive Antibodies Elicited by Pathogens: The Good, the Bad and the Ugly. Curr Med Chem. 2017 May 9;24(36).

173. Keasey SL, Schmid KE, Lee MS, Meegan J, Tomas P, Minto M, et al. Extensive antibody cross-reactivity among infectious gram-negative bacteria revealed by proteome microarray analysis. Molecular and Cellular Proteomics. 2009 May;8(5):924–35.

174. Damelang T, de Taeye SW, Rentenaar R, Roya-Kouchaki K, de Boer E, Derksen NIL, et al. The Influence of Human IgG Subclass and Allotype on Complement Activation. The Journal of Immunology. 2023 Dec 1;211(11):1725–35.

175. Jennewein MF, Alter G. The Immunoregulatory Roles of Antibody Glycosylation. Vol. 38, Trends in Immunology. Elsevier Ltd; 2017. p. 358–72.

176. Liao M, Ye F, Zhang B, Huang L, Xiao Q, Qin M, et al. Genome-wide association study identifies common variants at TNFRSF13B associated with IgG level in a healthy Chinese male population. Genes Immun. 2012;13(6):509–13.

177. Pollack M., Huang A. I., Prescott R. K., Young L. S., Hunter K. W., Cruess D. F., et al. Enhanced survival in Pseudomonas aeruginosa septicemia associated with high levels of circulating antibody to Escherichia coli endotoxin core. Journal of Clinical Investigation. 1983;72(6):1874–81.

178. Leitao Filho FS, Mattman A, Schellenberg R, Criner GJ, Woodruff P, Lazarus SC, et al. Serum IgG Levels and Risk of COPD Hospitalization: A Pooled Meta-analysis. Chest. 2020 Oct 1;158(4):1420–30.

